# Enabling single-molecule localization microscopy in turbid food emulsions

**DOI:** 10.1101/2021.03.03.433739

**Authors:** Abbas Jabermoradi, Suyeon Yang, Martijn Gobes, John P.M. van Duynhoven, Johannes Hohlbein

## Abstract

Turbidity poses a major challenge for the microscopic characterization of many food systems. In these systems, local mismatches in refractive indices can cause reflection, absorption and scattering of incoming as well as outgoing light leading to significant image deterioration along sample depth. To mitigate the issue of turbidity and to increase the achievable optical resolution, we combined adaptive optics (AO) with single-molecule localization microscopy (SMLM). Building on our previously published open hardware microscopy framework, the miCube, we first added a deformable mirror to the detection path. This element enables both the compensation of aberrations directly from single-molecule data and, by further modulating the emission wavefront, the introduction of various point spread functions (PSFs) to enable SMLM in three dimensions. We further added a top hat beam shaper to the excitation path to obtain an even illumination profile across the field of view (FOV). As a model system for a non-transparent food colloid in which imaging in depth is challenging, we designed an oil-in-water emulsion in which phosvitin, a ferric ion binding protein present in from egg yolk, resides at the oil water interface. We targeted phosvitin with fluorescently labelled primary antibodies and used PSF engineering to obtain 2D and 3D images of phosvitin covered oil droplets with sub 100 nm resolution. Droplets with radii as low as 200 nm can be discerned, which is beyond the range of conventional confocal light microscopy. Our data indicated that in the model emulsion phosvitin is homogeneously distributed at the oil-water interface. With the possibility to obtain super-resolved images in depth of nontransparent colloids, our work paves the way for localizing biomacromolecules at colloidal interfaces in heterogeneous food emulsions.

## Introduction

In turbid media, optical imaging can be compromised by the presence of ingredients or phases bearing different refractive indexes. In food emulsion such as mayonnaise, for example, the presence of oil droplets in the aqueous phase will disturb incoming and outgoing wavefronts of light. With an increasing depth of imaging, more and more photons will be reflected, absorbed, or scattered leading to aberrated images that suffer from decreased resolution, blurriness and distortions (Schwertner et al., 2004; Vellekoop and Aegerter, 2010). To correct aberrations, the concept of adaptive optics (AO) was developed in which active controllable elements such as deformable mirrors or spatial light modulators allow to modulate the wavefronts before the light reaches the photon detecting camera (Girkin et al., 2009). First developed for astronomical telescopes (Hardy, 1998), AO is increasingly finding applications in fluorescence microscopy such as a super resolution (Booth et al., 2015), light-sheet (Dalgarno et al., 2012), confocal (Hell et al., 1993; Ji et al., 2012) or multiphoton (Dong et al., 2003) microscopy. A common implementation of AO uses a deformable mirror in reflecting mode to compensate aberrations that can be described by Zernike polynomials (Madec, 2012). The deformable mirror consists of a number of tiltable micromirrors or actuators allowing to modulate individual sections of light in the Fourier plane. Deformable mirrors can be operated in an open loop or in a closed loop (Booth, 2014). In closed loop operation, an additional wavefront sensor (e.g., Shack-Hartmann wavefront sensor) is required to send instantaneous feedback to the deformable mirror (Tao et al., 2011). However, operation in closed loop reduces the number of photons available for image formation, hampering especially applications in super-resolution localization microscopy (SMLM) (Betzig et al., 2006; Hess et al., 2006). In SMLM, individual emitters, whose distance to each other is below the diffraction limit of optical microscopy, can be distinguished from each other, if conditions are achieved that allow separating the emission of each fluorophore in time. In the dSTORM (direct stochastic optical reconstruction microscopy) variant, this requirement is achieved by using blinking fluorophores that switch between fluorescent and non-fluorescent states (Heilemann et al., 2008; Rust et al., 2006). Originally a 2D imaging technique, 3D resolution in SMLM can be achieved by breaking the axial symmetry of the imaged point spread function (PSF) using astigmatism (Huang et al., 2008; Holtzer et al., 2007) or further PSF engineering via phase masks or AO enabling saddle point, tetrapod (Aristov et al., 2018) and double helix (Pavani et al., 2009) PSF. Our recent work introduced a method called circular-tangent phasor-based SMLM (ct-pSMLM) that enables fast and accurate localization of emitters after PSFs engineering on standard CPUs (Martens et al., 2020). For our purposes, we will use a sensorless approach based on a deformable mirror to compensate aberrations and introduce engineered PSFs providing access to three-dimensional super-resolution microscopy. In particular, we adapted an approach called REALM (Robust and Effective Adaptive Optics in Localization Microscopy) that was recently developed to compensate aberrations in depth of complex biological samples (Siemons et al., 2020). REALM uses the image quality metric derived from a weighted sum of Fourier transforms of raw images of emitters to estimate the aberrations. REALM then compensates the aberrations of different Zernike modes based on the metric values and biases of the mirror. Here, to establish SMLM as a powerful method to study food related turbid media, we updated the miCube microscopy framework (Martens et al., 2019) on several aspects. Quantitative SMLM measurements are often compromised by inhomogeneous illumination due to a Gaussian intensity distribution of the exciting laser beam. To overcome this issue, several approaches have been introduced to achieve illumination with a constant intensity over the entire field of view. Examples include the use of a pair of micro lenses array consisting of identical spherical lenslets (Douglass et al., 2016; Scholtens et al., 2011), the use of multimode optical fibers for illumination (MMF) (Deschamps et al., 2016) in combination with speckle reducers (Kwakwa et al., 2016) or rotating diffusers (Ma et al., 2017) with the latter being less suitable for total internal reflection fluorescence (TIRF) microscopy due to the degradation of spatial coherence prevention diffraction limited focusing. Recent work further demonstrated flat field illumination over variable field sizes using two galvanometer scanning mirrors placed in a plane conjugated to the back focal plane of the microscope objective in epifluorescence or TIRF mode (Mau et al., 2020). In our implementation, we added a top-hat beam shaper in the excitation path that converts the Gaussian shaped intensity distribution of the excitation beam into a homogeneous flat field profile (top hat) enabling quantitative microscopy (Khaw et al., 2018; Stehr et al., 2019). Moreover, we equipped the miCube with a deformable mirror placed in the detection pathway to compensate aberrations coming from in-depth imaging of opaque samples with spatial variations of refractive indices and for enabling PSF engineering. As a model for mayonnaise, a highly turbid food emulsion containing up to 80% of oil, in which egg yolk acts as an emulsifier (Depree and Savage, 2001), we will demonstrate our approach on a dilute oil-in-water model emulsion that was emulsified with phosvitin (Castellani et al., 2006). Phosvitin is a protein contained in egg yolk that has a binding capacity for ferric ions (Zhang et al., 2011). Ferric/ferrous ions can catalyze lipid oxidation at the oil-water interface, which can detriment the sensorial and nutritional quality of food emulsions. Visualization of phosvitin at oil-water interfaces in food emulsions is therefore relevant to understand lipid oxidation mechanisms and designing anti-oxidant strategies (Berton-Carabin et al., 2014). For our model emulsion, we opted for a 15% v/v oil concentration to obtain small droplets (~ 1 μm diameter) with a high surface area available for phosvitin. We then used a phosvitin antibody conjugated with the fluorophore Alexa Fluor 647 to map phosvitin at the droplet interface.

## Materials and Methods

### The miCube excitation path

For laser excitation, we used one of two options. The first option features a standard laser combiner (Omicron Lighthub, Germany) equipped with 4 lasers operating at 405 nm (60mW), 488nm (200mW), 561 nm (500mW), and 642nm (2x 200mW) that were coupled into a singlemode fiber. As a second option, we used two low-cost diode lasers equipped with simple beam shaping optics. The lasers operate at 635 nm (PD-01287, 200 mW, Standard Module, Lasertack, Germany) and 520 nm (PD-01298, 100 mW, Standard Module, Lasertack, Germany) and are controlled via a home-built Arduino powered laser control engine (hohlbein-lab.github.io/miCube/LaserTrack_Arduino.html). After combining the laser light with a dichroic mirror (RGB Beam Splitter-Combiner, Lasertack, Germany), the light was coupled into a single-mode fiber (P3-460B-FC-2, Thorlabs) using a 10x objective (RMS10X - 10X Olympus Plan Achromat Objective, Japan). The coupling efficiencies of the 635 nm and 520 nm diode lasers to single-mode fiber were 56% and 42% respectively. For collimating the diverging laser light after the fiber, we used an achromatic lens (CL) of either 30mm or 60mm focal length (AC254-030-A-ML and AC254-060-A-ML, Thorlabs), and ensured with a shear plate (SI050, Thorlabs) proper collimation of the laser light. The parallel beam was then reflected by an elliptical mirror (M1, BBE1-E02, Thorlabs) mounted to a right-angle kinematic cage mount with ±4° pitch and yaw adjustment (KCB1E/M, Thorlabs), towards the top-hat beam shaper (TSM25-10-D-D-355, Top Shape, Asphericon GmbH, Germany) to create a homogeneous distribution illumination intensity. The ideal input beam size for the beam shaper is between 9.2mm and 10.8mm (1/*e*^2^). Using the CL = 60 mm, the input beam size is approximately 10.2 mm. After the beam shaping, the laser beam was reflected with another mirror (M2, BBE1-E02, Thorlabs) towards an iris (Iris, SM1D12D, Thorlabs) to control the illumination area in the sample plane. The combination of M1 and M2 was used to position the beam in respect to the top-hat beam shaper. The laser light was then focused into the back focal plane (BFP) of the microscope objective using an achromatic lens (TR, f = 150mm, AC508-150-A-ML, Thorlabs) mounted on a translational stage (XR25C/M, Thorlabs) used to change the position of the focus in the back focal plane.

### The miCube main block

The main block itself is similar to the one reported previously (Martens et al., 2019) and connects the excitation and emission paths with the sample. The cube is made from aluminum and anodized in black to suppress stray light. The block has three openings for the light to pass, two at faced sides for the emission and the detection path and one at the top threaded (M25 x 0.75) such that the microscope objective can be directly screwed in. The laser beam focused by the TIRF lens is reflected by a polychroic mirror (DiM, ZT532/640rpc or ZT405/488/561/640rpcv2, Chroma) into the back focal plane of the objective lens (OL, CFI Plan Apo Lambda 100x Oil NA 1.45, Nikon). The sample was placed on 3D printed coverslip sample holder and secured in place with small magnets. We used a stick-slip piezo XYZ stage (SLS −3232, SmarAct GmbH, Germany) for sample scanning. The stage has a footprint of 32 mm by 32 mm and offers a travel range of 21 mm in each direction with 1 nm closed-loop resolvable position resolution. The stage able to handle payloads of up to 1.5 N. The light emitted from the sample was collected with the same microscope objective and, after passing the polychroic mirror, further cleaned up with a bandpass filter (F, ZET532/640m-TRF, Chroma) located at the bottom of the dichroic cage holder (DFM1/M, Thorlabs) to block remaining back-reflected laser light from entering the emission path. Subsequently, emitted light was reflected using a 90-degree mirror (M3, BBE1-E02, Thorlabs) and conducted to out of the cube and towards the tube lens.

### The miCube emission path

A tube lens (TuL, MXA20696, Nikon) with 200 mm effective focal length was implemented to focus the collimated light from the infinity-corrected objective into a first virtual imaging plane. The tube lens was placed in a 3D printed enclosure which was directly mounted to the main block. The focused light was reflected by an elliptical mirror (M4, BBE1-E02, Thorlabs) and conducted towards a 4f system of lenses. The first lens (L1, AC508-100-A-ML, Thorlabs) was positioned to collimate the light from the tube lens. For modifying the incoming wave front and to compensate the aberrations, we put the deformable mirror (DM, DMP40/M- P01, 40-Actuator Piezo Deformable Mirror, Thorlabs) in the Fourier plane of L1 (one focal distance). The deformable mirror consists of a 40-actuator array with 3 bimorph benders for ±2.0 mrad Tip/Tilt actuation and was mounted on a XZ linear stage (XR25C/M, Thorlabs) to simplify the alignment of the mirror in respect to the emission light. Moreover, as the angle between incident and reflected light from the deformable mirror should stay below 30 degrees, we used another mirror (M5, PF10-03-P01, Protected Silver Mirror, Thorlabs) mounted to a precision kinematic mirror holder (KS1, Thorlabs), placed in front of the deformable mirror to control this angle. The light reflected from the deformable mirror was conducted to the second lens (L2, AC508-100-A-ML, Thorlabs) of the 4f system, which focused the light on the camera (Prime 95B sCMOS, Photometrics) having a maximum quantum yield of 95% QE and featuring a 11 μm by 11 μm pixel size. The camera was mounted on a custom 3D printed stand to adjust the height and position on the optical table.

### Re-Scan Confocal Microscopy

We updated the previous miCube microscope with a rescan confocal microscopy (RCM) module (Confocal.nl, Amsterdam, The Netherlands) (Luca et al., 2013). The RCM has a separate fiber input for and allows scanning the collimated laser beam across the sample. The emitted light from the specimen is then rescanned with a second mirror with twice the sweep length as the excitation scanning mirror, leading to a 43 nm pixel size on the sCMOS chip. In practice, RCM is capable of a 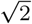 times increase in resolution compared to classical confocal laser scanning microscopes.

### Deformable mirror for adaptive optics and PSF engineering

The mirror (DM, DMP40/M-P01, 40-Actuator Piezo Deformable Mirror, Thorlabs) consists of 40 individual actuators and three arms for tip/tilt. Each actuator and each arm can be controlled by applying voltages between 0 V and 200 V. In combination, the voltages determine the curvature of the mirror. To correct the flatness of the mirror and later for modulating the PSFs, we implemented REALM (github.com/MSiemons/REALM) (Siemons et al., 2020), which allows for corrections without requiring an additional sensor to monitor the incoming wavefront. We further wrote a plugin for Micromanager to connect REALM to the deformable mirror used in our study (github.com/HohlbeinLab/Thorlabs_DM_Device_Adapter) Martens et al. (2020).

### Visualization and image analysis of SMLM data

The raw data were analyzed with ThunderSTORM (Ovesný et al., 2014) plugin in ImageJ/Fiji (Schindelin et al., 2012) based on our phasor-based localization algorithm (Martens et al., 2019). After obtaining the localizations, we performed two-dimensional cross-correlation drift correction (settings in ThunderSTORM: 10 bins and 5x magnification). The localizations were visualised using the average shifted histogram options, with the magnification set to 5. Moreover, no lateral shifts were added and cyan was chosen for the lookup table. A rewritten ImageJ plugin was used to remove constant fluorescence background by means of a temporal median filter (Nieuwenhuizen et al., 2013) (see github.com/HohlbeinLab/FTM2 for the ImageJ/Fiji plugin). To determine the histogram of droplet sizes, we first manually encircled droplets in the field of view based on the ring-shaped presence and absence of fluorescence. We then applied the Hough circle transform (Yuen et al., 1989) function in MATLAB (Mathworks, UK) to obtain the radii of all circles using 0.2 μm as the minimum and 2 μm as the maximum search radius for oil droplets. For measuring the resolution of super resolved images, we further used Fourier Ring Correlation (FRC) as implemented in the software package SMAP (Ries, 2020).

### Isolation and purification of phosvitin

Our procedure of isolating and purifying phosvitin follows previous work (Zhang et al., 2011). Briefly, fresh hen eggs were purchased from the domestic market. To remove egg white and chalazas, the yolks were rolled on a filter paper. The temperature of the following steps was kept at 4°C. An equal amount of distilled water was added to the yolk. The diluted solution was centrifuged at 12000g for 10min (Avanti j-25, Beckman). The precipitate was collected and homogenized with an equal mass of a 0.17M NaCl solution and centrifuged again at 12000 g for 10 min. The granules were dissolved in a 10% w/v of a 1.74M NaCl solution. The pH was adjusted to 8.0 withlmM NaOH solution and homogenized with 4% PEG6000 w/w and centrifuged at 12000 g for another 10 min. The supernatant was dialyzed against distilled water for 48 hours and subsequently centrifuged at 12000 g for 10 min. The supernatant was collected and lyophilized using a lyophilizer from either (Christ, Germany) or (Labconco, USA).

### Phosvitin based model emulsion

6mgmL^−1^ of lyophilized phosvitin was added to 0.5 M of 2-N-porpholino ethane sulfonic acid (MES) buffer at pH 6.6. The solution was centrifuged at 4000 g for 20 min and the supernatant was extracted to remove the undissolved particle from the solution. We then added 0.15% w/v of sodium dodecyl sulfate (SDS) to the solution to obtain a stable model emulsion. A 15% oil in water mixture was prepared with 7.5 mL of rapeseed oil and 42.5 mL of the phosvitin containing solution. The emulsion was premixed with a 18 mm diameter head disperser at 18000 rpm for 2 min (T 18 digital ULTRA-TURRAX, IKA, Germany). Next, the 15% oil in water model emulsion was obtained by emulsifying the premix at 70 bar for 20 min with a flow rate of 80 mL/ min using a high-pressure homogenizer (Delta Instruments LAB Homogenizer).

### Sample preparation

For dSTORM measurements, the phosvitin antibody conjugated with Alexa 647 stock solution was first diluted 50 times in TRIS buffer. 40 μL of the diluted antibodies were then added to 400 μL of the 15% oil-in-water model emulsion. In order to stall self-diffusion of droplets, we further added 0.5% w/v of guar gum (Sigma, ref. G4129). The specimen was then dripped into a well of a silicon gasket. 1.5 μL of the STORM buffer containing 50 mM TRIS pH8, 10 mM NaCl, 10% glucose, 140 mM 2-mercaptoethanol, 68 pgmL^−1^ catalase, and 200μgmL^−1^ glucose oxidase (Jimenez et al., 2020) was mixed into 15 μL of sample, before we sealed the gasket. We note that although we effectively used a tenfold reduced concentration of a standard STORM buffer in our sample, sufficient blinking was achieved. We tried using a 10x concentration of BME which did not lead to improvements in blinking and achievable resolution. For the confocal laser scanning microscopy (CLSM) measurements on mayonnaise, 1% w/w of 1mg mL^−1^ Nile blue (Sigma, ref.N0766) solution was gently stirred into the mayonnaise. For CLSM experiments on the model emulsion, we further added 0.5% w/v guar gum to prevent droplet self-diffusion. For correcting the deformable mirror, we used fluorescent beads (FluoSpheres™ Carboxylate-Modified Microspheres, 28 nm diameter, dark red fluorescent (660/680), Thermo Fisher). First, we diluted the provided solution 1:100000 and added 4 μL of the dilution to a coverslip (#1.5, Thermo Scientific Menzel Gläser). We then used a second coverslip used on top of the first one to have homogeneous distribution of beads on the field of view. To measure the drift characteristics in x, y and z, we prepared a sample as described but using 50 nm fluorescent beads (560 nm peak emission wavelength) instead.

## Results and Discussion

### Turbidity limits the optical resolution and in-depth imaging of oil-in-water emulsions

To demonstrate the challenges of optical imaging in turbid media, we first imaged a mayonnaise sample using a stack of confocal images (Figure 1A). We added Nile blue to the mayonnaise and excited at 642 nm. The expected loss of optical resolution in depth of the sample is seen in the cross-sectional view; an increase of imaging depth coincides with increasing blurriness (Figure 1B,C).

**Fig. 1.**
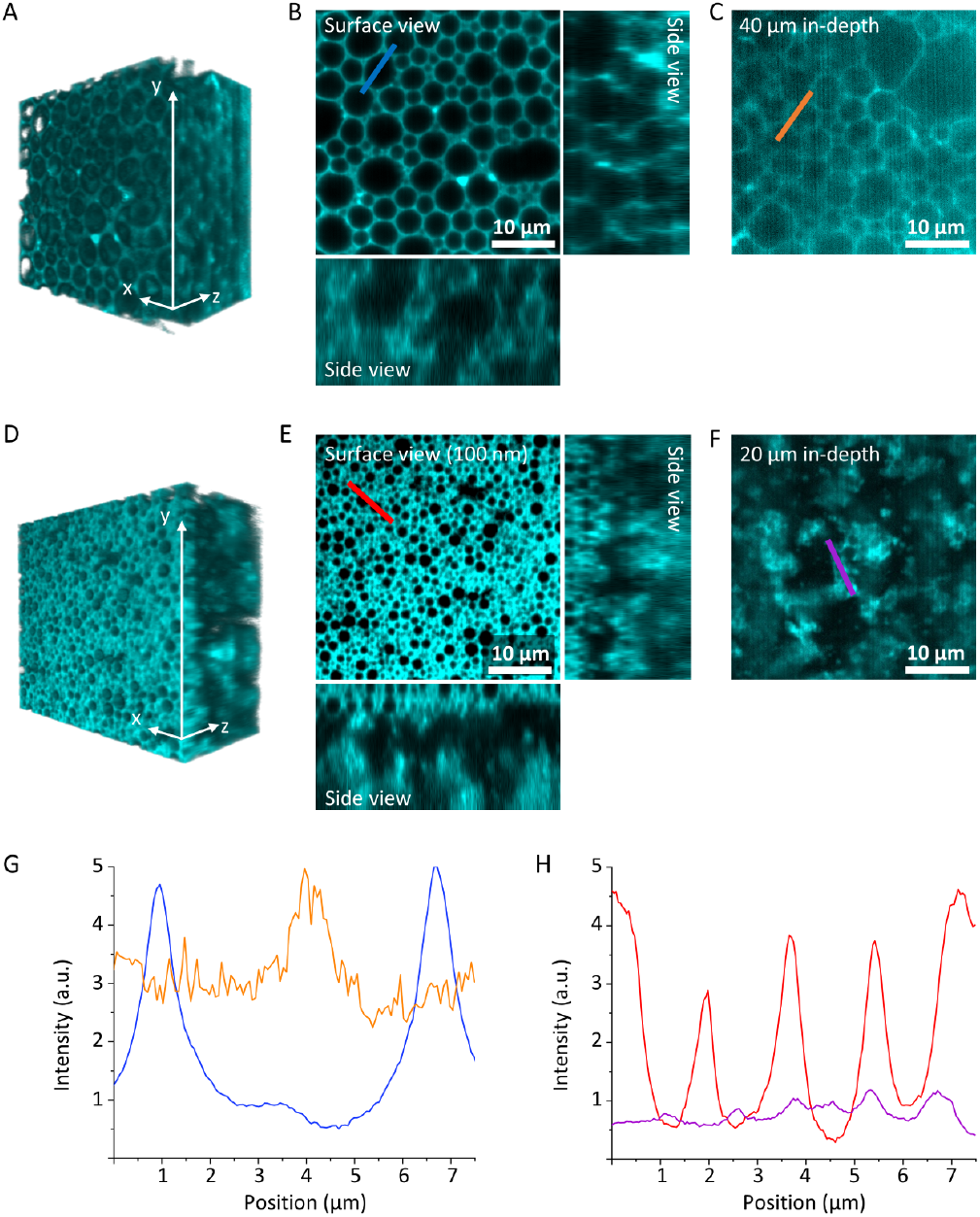
Re-scan confocal images of mayonnaise and the model emulsion labeled with Nile blue and excited at 642 nm. **A)** 3D volume rendering image of 40 μm by 40 μm by 40 μm taken with a 1 μm step size along the optical z-axis. **B,C)** Cross-sectional views in different planes of xy-xz-yz shows in a slice of 20 μm thickness (B) and in the xy plane located in 40 μm depth in the sample (C). **D)** 3D volume rendering of 40 μm by 40 μm by 20 μm stabilized by guar gum, taken with 1 μm step size. **E,F)** Cross-sectional views in different planes of xy-xz-yz starting at the surface close to the coverslip (E) and in the xy plane located in 20 μm depth in the sample (F). **G)** Line plots of fluorescence intensity (blue line in B) representing the achievable signal to noise of droplets close to the glass interface and (orange line in C) in 40 μm depth. **H)** Line plots of fluorescence intensity (red line in E) representing the achievable signal to noise of droplets close to the glass interface and (purple line in F) in 20 μm depth.

### Developing a model system for turbid oil-in-water emulsions

To obtain a model emulsion suitable for SMLM, we prepared a low (15%) oil-in-water model emulsion emulsified with phosvitin. Due to the lower oil content of the model emulsion, the packing of droplets is less dense leading to significant self-diffusion of droplets. We therefore added guar gum to the emulsion which increased the viscosity leading to an effective immobilization of the droplets. The rescan-confocal laser scanning images showed that the high shear during emulsification resulted in smaller droplet sizes (~ 0.5 μm radius) compared to typical mayonnaises (2-2.5 μm radius) (Figures 1D). Similar to imaging in mayonnaise, we observed a loss of resolution and signal-to-noise ratio with increasing depth (Figure 1). We note that larger areas are void of droplets and are likely occupied with guar gum networks (Figure 1E,F: xz and xy sectional views).

### Adapting the miCube for super-resolution localization mi-croscopy in turbid media

We modified the miCube microscopy framework (Martens et al., 2019) at various positions to address the challenges imposed by super-resolution measurements in turbid media and to provide users with additional hardware options (Figure 2). In the excitation path, we included an option to use cheaper laser diodes as light sources rather than a scientific grade multicolor laser engine. To enable simplified quantitative analysis of super-resolution data, we added a top-hat beam shaper providing an even illumination profile over the field of view (Khaw et al., 2018; Stehr et al., 2019; Douglass et al., 2016). For the main cube, we opted for a sample scanning stage, offering nanometer resolution over an 18 mm scanning range in all three directions and working in closed loop mode to compensate the thermal drift of the stage (Supplementary Figure 1). In the detection path, we implemented a deformable mirror as adaptive optics to correct the aberrations and enable PSF engineering that allow us to have higher z range in 3D. Moreover, we added a sCMOS camera with 95% quantum efficiency.

**Fig. 2.**
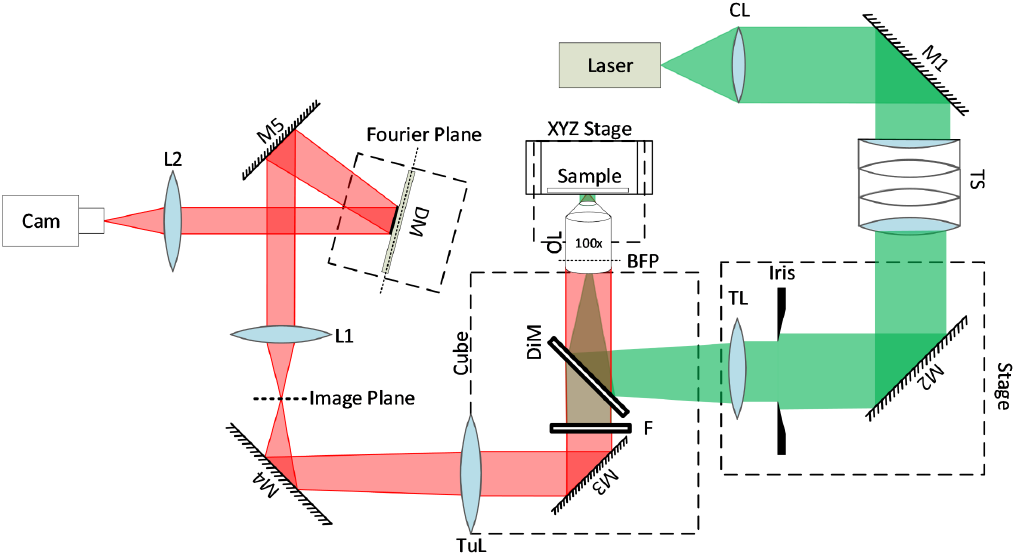
Schematic of the optical pathway. The excitation path (green) and the detection path (red) are highlighted. Components include a collimating lens (CL), mirrors (M), top-hat beam shaper (TS), TIRF lens (TL), polychroic mirror (DiM), objective lens (OL), back focal plane (BFP), bandpass filter (F), tube lens (TuL), lenses (L), deformable mirror (DM), and a camera (Cam).

### A flat top beam shaper provides an even illumination profile over a large field of view

We first inserted a flat top beam shaper in the excitation path (Figure 3) and compared the resulting illumination profile in the field of view with one obtained using conventional Gaussian distributed laser excitation. To this end, we placed a droplet of 1 μM Cy3B dye solution (Cytiva) on the coverslip and used a second coverslip on top to obtain a homogeneous spatial distribution of fluorophores. The fluorescence intensity profiles of the Gaussian and flat-top epi-illumination were obtained by exciting the sample with the 640 nm laser set to 26 mW. Using a collimating lens with a focal length of CL = 30 mm and 60 mm (for positioning see Figure 3), we achieved a full width at 90% of the maximum intensity (FW90M) for these two lenses in the Gaussian illumination mode of 27 μm and 28 μm and for flat illumination of 116 μm and 128 μm, respectively (Figure 3C). The flat field leads to laser intensity of 0.19kW cm^−2^.

**Fig. 3.**
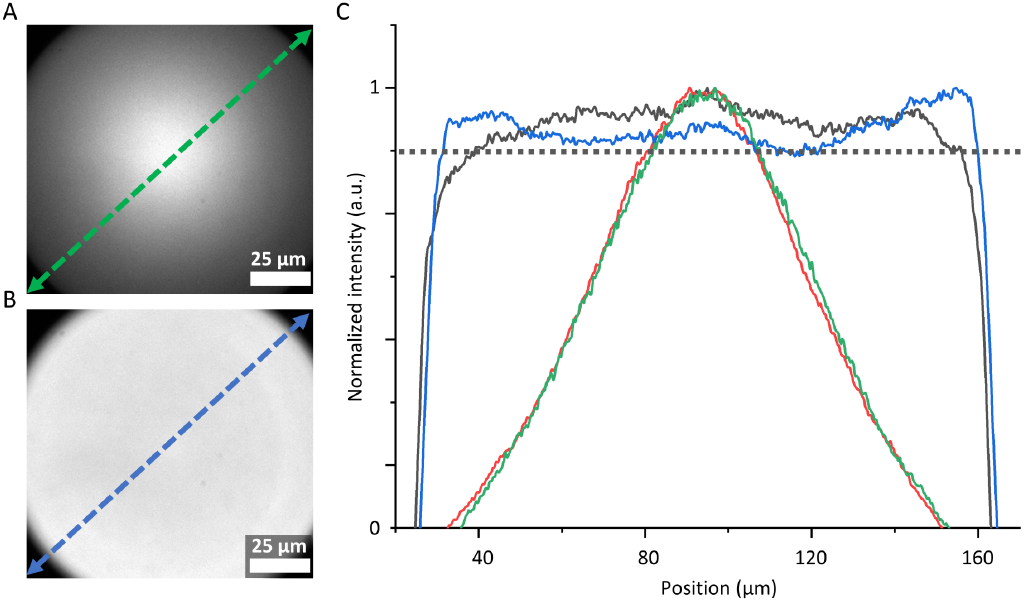
Characterization of the intensity profile in the field of view. Two collimating lenses were compared after the fiber output with either 30 mm and 60 mm focal length with and without an addition top-hat beam shaper present in the collimated laser beam. **A)** Gaussian beam profile after using the 60 mm lens for colimitation. **B)** 60 mm lens with added top-hat beam shaper. **C)** The line profiles of A, green line) and (B, blue line) are plotted and compared to using the 30 mm lens for collimation in absence (red line) and presence of the top-hat beam shaper.

### Deformable mirror correction and characterization

We implemented a deformable mirror to compensate for aberrations introduced by either the sample or by other optical elements in the detection pathway. We noted, however, that the deformable mirror itself introduces additional aberrations to the system requiring corrections. Using the standard setting of all actuators set to 100 V, we observed asymmetrically elongated PSFs (Figure 4A) rather than the expected symmetrical and circular PSFs when imaging fluorescent latex beads of 28 nm diameter. To correct the flatness of the mirror and later for modulating the PSFs, we implemented REALM (Siemons et al., 2020). Using REALM, each Zernike aberration mode was individually corrected by sequentially optimizing the image metric (Figure 4B). The software evaluated 11 biases ranging from —100 nm to 100 nm for the mirror setting for each of the 9 tested Zernike modes. By using 3 correction rounds, a total of 297 images was acquired. A Gaussian function was fitted to the metric values of each bias. The position of the maximum of the fitted function was taken as the required correction amplitude for that specific mode. After correcting the mirror, the expected circular symmetry of the PSF is restored (Figure 4c bottom) and we found the correction settings to be stable for several months.

**Fig. 4.**
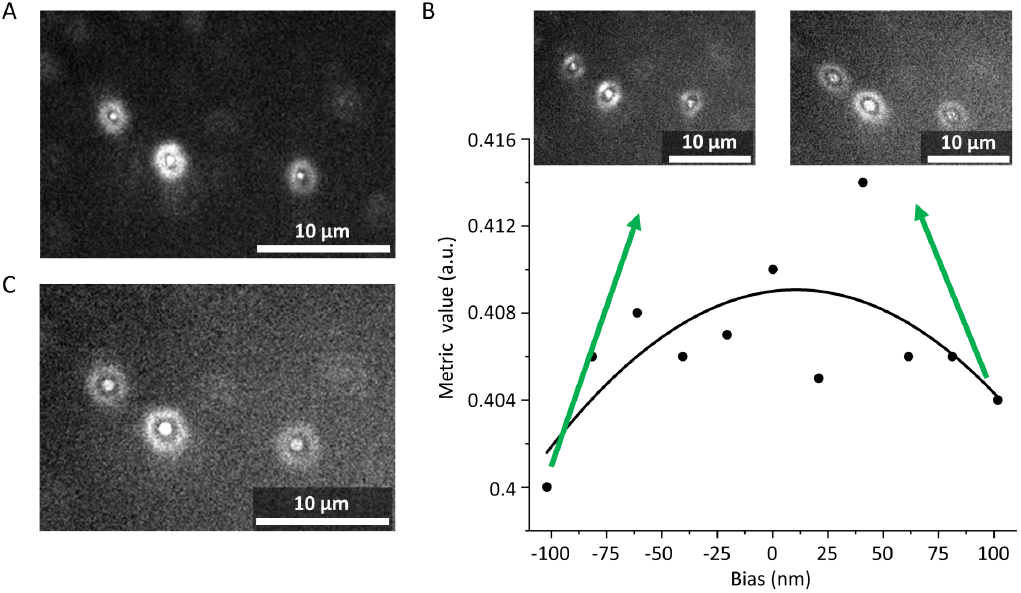
Corrections of aberrations using adaptive optics. **A)** Fluorescent beads (28 nm diameter) were immobilized on a cover slip and imaged. Here, the asymmetrical shape of the PSF is induced by the deformable mirror itself. **B)** A specific Zernike mode, oblique secondary astigmatism -Z(4,-2), was selected and the correction procedure was performed providing a Gaussian fit of the obtained metric values and biases. **C)** The expected symmetrical shape of the PSF is restored after correcting the deformable mirror for all Zernike modes using REALM. We note that the beads are slightly out of focus to exemplify the symmetry.

### Super-resolution microscopy in turbid media

Phosvitin is an egg yolk protein that has a binding capacity for ferric ions (Zhang et al., 2011) which can catalyze lipid oxidation at oil-water interface in food emulsions. Here, we aimed at visualizing phosvitin at the oil-water interfaces of a model emulsion to explore the potential spatial heterogeneity of phosvitin that could provide clues to design strategies combating lipid oxidation. In our 15% oil-in-water model emulsion, phosvitin took the role as the main emulsifier. We further added SDS to increase the stability of the emulsion (Figure 5). To localize phosvitin at the oil-water interface, we added fluorescently labelled antibodies against phosvitin.

**Fig. 5.**
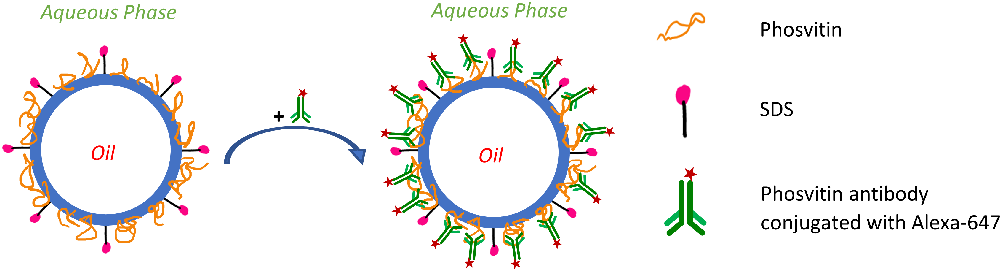
Schematic diagram of an oil in water emulsion droplet in the phosvitin stabilized model emulsion. Phosvitin and SDS jointly stabilize the oil-water interface. We then added antibodies against phosvitin that are fluorescently labelled (Alexa 647) to localize phosvitin at the interface using dSTORM.

After targeting the phosvitin with the antibody, we imaged the specimen in a plane close to the surface (a few 100 nm deep) to minimize aberrations. Using the 640 nm laser for excitation and 405 nm for photo-reactivation of the Alexa-647 fluorophores we recorded a video of 10000 frames with 20 ms frame time. A rewritten temporal median filter (Nieuwenhuizen et al., 2013) was used to remove constant fluorescence background before the positions were obtained by our phasor-based localization approach. Figure 6A shows the outlines of the oil droplets representing labelled antibodies bound to phosvitin. To verify the specificity of the phosvitin antibody, a control experiment was performed with a phalloidin antibody conjugated with Alexa-647. As expected, a noisy background was observed without clear outlines of droplets being visible (Supplementary Figure 2). For the phosvitin measurements close to the glass surface, we plotted a line profile over two isolated droplets (Figure 6a) indicating two radii of ~ 0.65 μm and ~ 0.75 μm, respectively. Further, analyzing the profile of the droplets ring using a Gaussian fit and calculating the FWHM of the intensity profile around the droplets revealed a thickness of 74 nm (Supplementary Figure 3) thereby representing a convolution of the localization precision and the expected geometrical averaging due to projecting a three-dimensional cut out of a sphere onto a two-dimensional imaging plane with. Moreover, we obtained an imaging resolution of 71 nm using FRC (Nieuwenhuizen et al., 2013; Ries, 2020; Banterle et al., 2013) (Supplementary Figure 4).

**Fig. 6.**
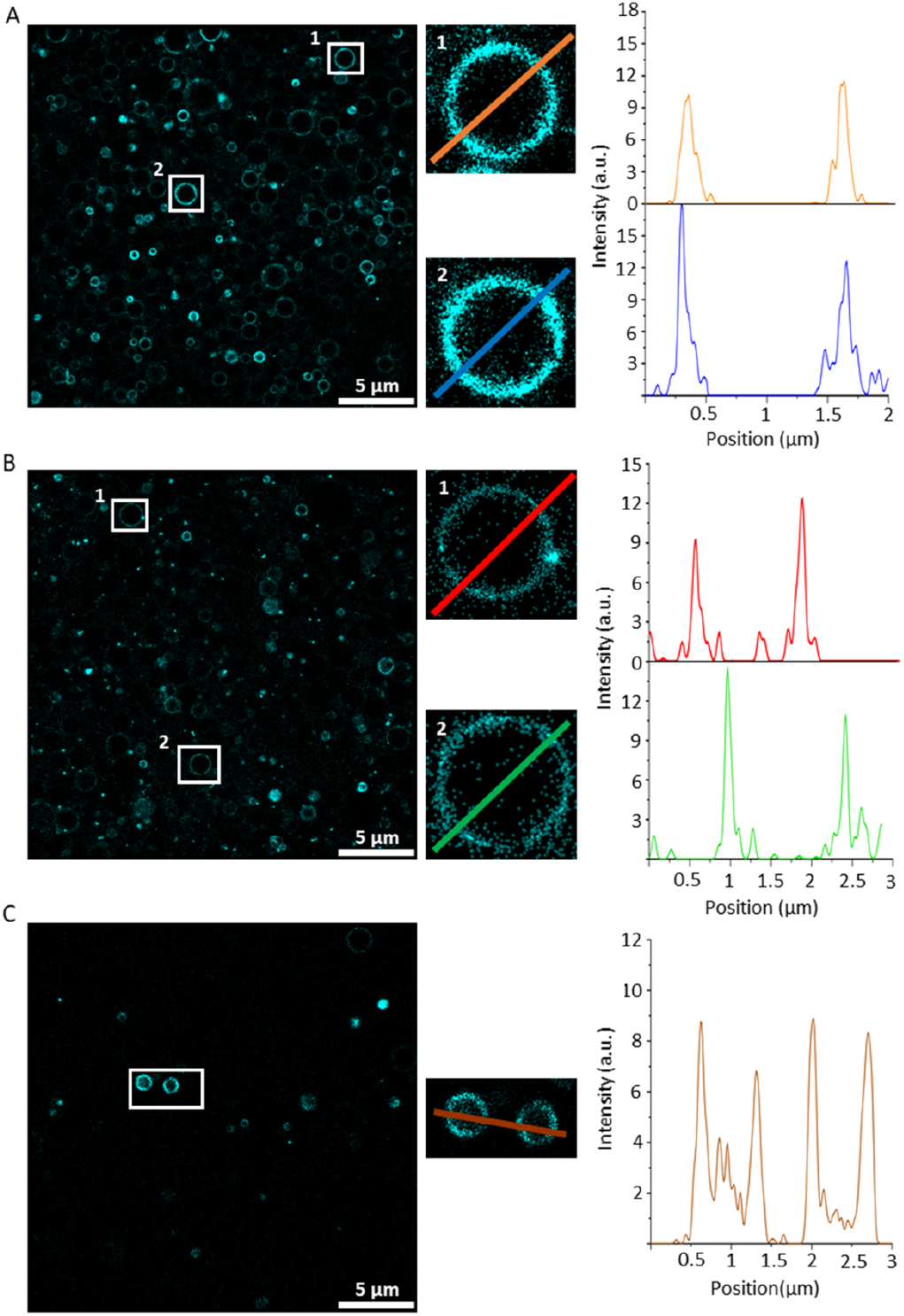
Super resolved imaging in an oil-in-water model emulsion in which fluorescently labelled antibodies are bound to phosvitin present at the oil-water interface. **A)** 25 μm by 25 μm image of the super-resolved droplets close (<1 μm) to the surface. The right images show two enlarged droplets from the sample with a radius ~ 0.65 μm (top) and ~ 0.75 μm (bottom) as obtained in the line profile. **B)** 25 μm by 25 μm image of droplets in 4 μm depth. The two enlarged droplets have a radius of ~ 0.75 μm (top) and ~ 0.70 μm (bottom). **C)** Image of droplets in 15 μm depth. The two enlarged droplets both have a radius of ~ 0.45 pm.

In a second experiment, we recorded 10000 frames in 4 μm depth (Figure 6B) after applying the aberration corrections obtained from single emitters with REALM. In this depth, the PSFs were only slightly aberrated such that the emission from individual fluorophore emitters could be directly used to obtain the correction coefficients. We again enlarged two droplets and obtained a radius of 0.75 μm and 0.7 μm, respectively. In this plane, we obtained 79 nm for the FWHM of the intensity profile around the droplets (Supplementary Figure 3). In a third experiments, we imaged 15 μm in depth after having to correct the PSFs using with embedded fluorescent latex beads. In this depth, we had to increase the laser excitation power twofold to obtain a sufficient number of photons per localization (Figure 6C). As we measured deeper into the sample, we noticed a decrease in the number of droplets present likely induced by the guar gum used reduce the mobility of droplets in the sample. We achieved super resolved images in 15 μm depth indicated by resolving two droplets with ~ 0.45 μm radius. Using FRC, we calculated the resolution of the image to be ~ 124nm (Supplementary Figure 4). For our measurement in the plane close to the surface, we found 284 droplets with radii between 0.2 μm to 1.2 μm with a number averaged mean radius of 0.46 μm (Figure 7A and Supplementary Figure 5). We note that these are apparent radii due to the error introduced by the imaging plane not crossing droplets at the center (Dalen and Koster, 2012; Schuster et al., 2012). For the images in 4pm depth, we counted 134 droplets with a different radius between 0.2 μm to 1.8 μm and a number averaged mean of 0.55 μm. As expected, when measuring deeper 15 μm depth) in the sample, the number of droplets per field of view reduced to 27 with a number averaged mean radius of 0.43 μm and mostly those droplets distributed between 0.35 μm and 0.48 μm. We note that the operational range offered by SMLM is not accessible by conventional CLSM, which typically has a lower limit of 0.5 μm for determining the radii of droplets (Duynhoven et al., 2002).

**Fig. 7.**
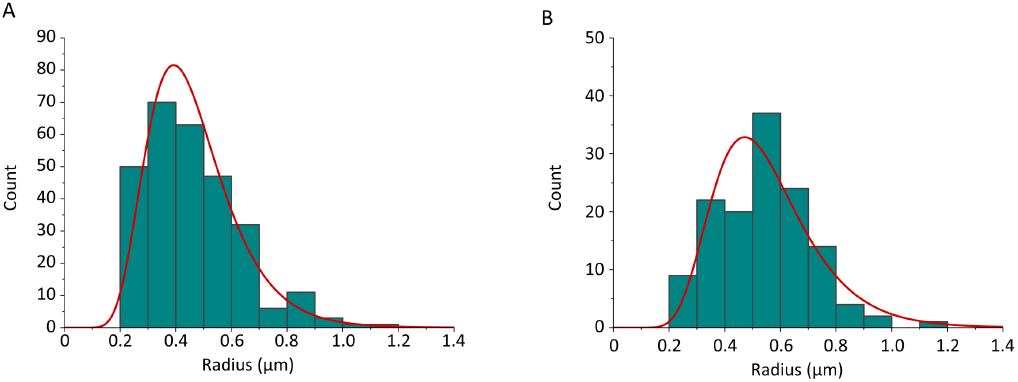
Apparent size distributions presented as histograms for a 15% model emulsion stabilized with phosvitin for **A)** the plane close to the surface and, **B)** the plane 4 μm in depth.

### Obtaining 3D image of oil droplets using PSF engineering

To show the capability of 3D imaging of adaptive optics, we recorded engineered PSFs using the deformable mirror. To access a 2.1 μm z range we employed vertical astigmatism and vertical secondary astigmatism Zernike (Saddle point PSF). We recorded 40000 frames with 30 ms frame time (Figure 8). We visualized the cross-sectional view of 3D distribution in xz and yz planes, showing that the full volume of a droplet can be covered and further indicating that phosvitin is homogeneously distributed at oil droplet interfaces. Vertical and horizontal dashed lines in Figure 8 indicate the corresponding xz and yz sections.

**Fig. 8.**
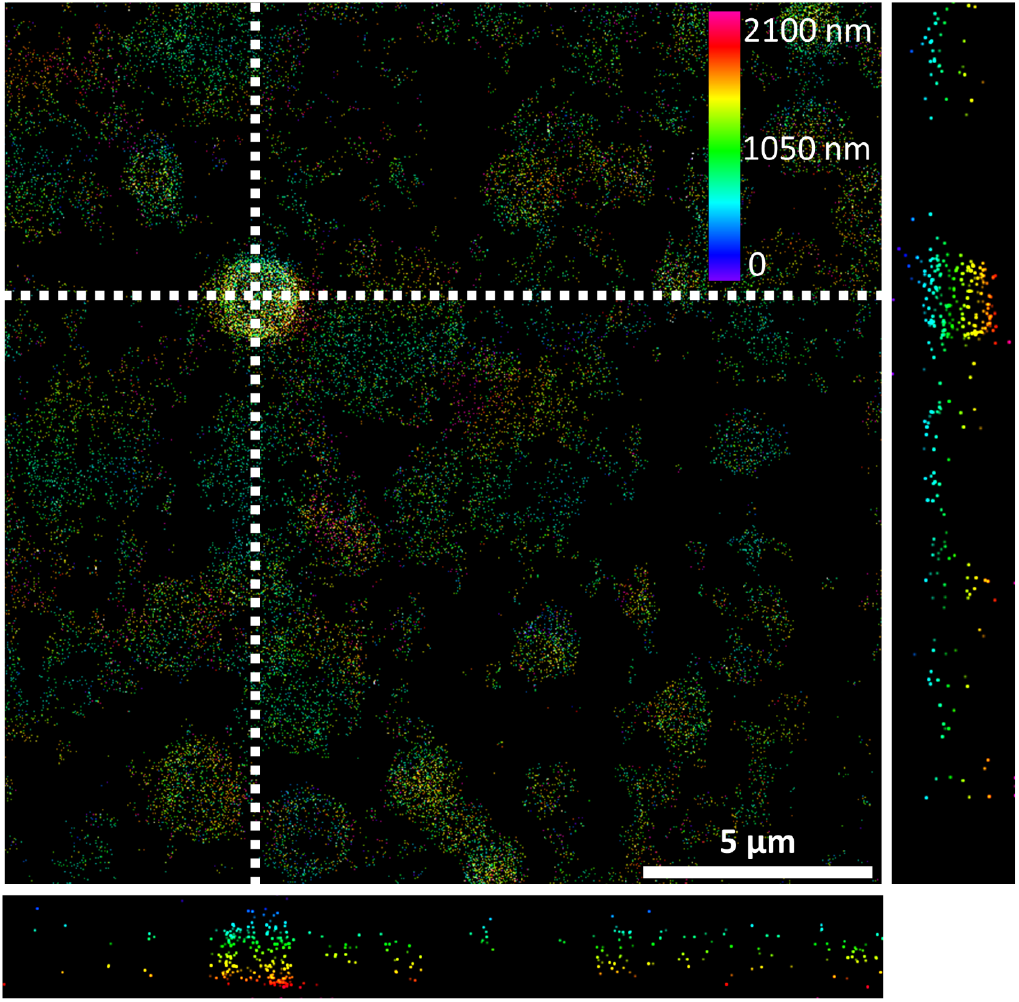
20 μm x 20 μm image of the oil in water model emulsion droplets using saddle point setting of the deformable mirror within a 2.1 μm z range. The side images show the distribution of protein in xz and xy. Saddle point was obtained by applying vertical astigmatism and vertical secondary astigmatism Zernike modes.

## Conclusion

In this study we presented an updated design of the miCube open-source microscope featuring flat field illumination and adaptive optics for PSF engineering. Together these updates enable 3D-SMLM in both standard samples and samples compromised by inherent turbidity. As a model system we used a dilute oil in water emulsion in which we imaged the iron-binding protein phosvitin at the droplet interface using a primary phosvitin antibody conjugated with Alexa 647. Flat field illumination enables homogeneous excitation intensities over areas of surpassing 100 μm by 100 μm indicating that phosvitin is homogeneously distributed over droplet interface. Droplets with radii as low as 0.2 μm can be discerned and localization of phosvitin in extended depths is possible. Using the the deformable mirror to engineer PSFs, we demonstrated that extended z-ranges can be accessed with SMLM without moving the focus of the objective in the sample plane. Our work showed the ability of the open hardware miCube platform to perform SMLM techniques for localizing biomacromolecules in both 2D and 3D at colloidal interfaces in complex food emulsions.

## Authors’ Contributions

A.J.: Methodology, Software, Validation, Investigation, Data curation, Drafting the manuscript, Writing—review and editing, Visualization. S.Y.: Investigation, Data curation. M.G.: Software. J.P.M.v.D.: Conceptualization, Methodology, Validation, Writing—review and editing, Supervision, Funding acquisition. J.H.: Conceptualization, Methodology, Validation, Writing—review and editing, Supervision, Funding acquisition.

## Data availability

The experimental raw data is currently available upon request and will be made available on https://zenodo.org.

## Acknowledgements

We would like to thank all our colleagues in the LocalBioFood consortium for helpful discussions.

## Supplement

**Supplementary Figure 1.** To measure the drift characteristics, we recorded 10000 frames of 50 ms each using 50 nm diameter beads excited at 561 nm. We engineered the PSF to represent a saddle-point PSF by changing the Zernike modes Z(2,-2) and Z(4,2) and recorded 3D raw data for determining the drift in the x, y, and z direction. We analyzed the data and performed 3D drift-correction using the cross-correlation function in SMALL LABS. Our data show that we have less than 200 nm drift in the lateral plane during the measurement.

**SFig 1.**
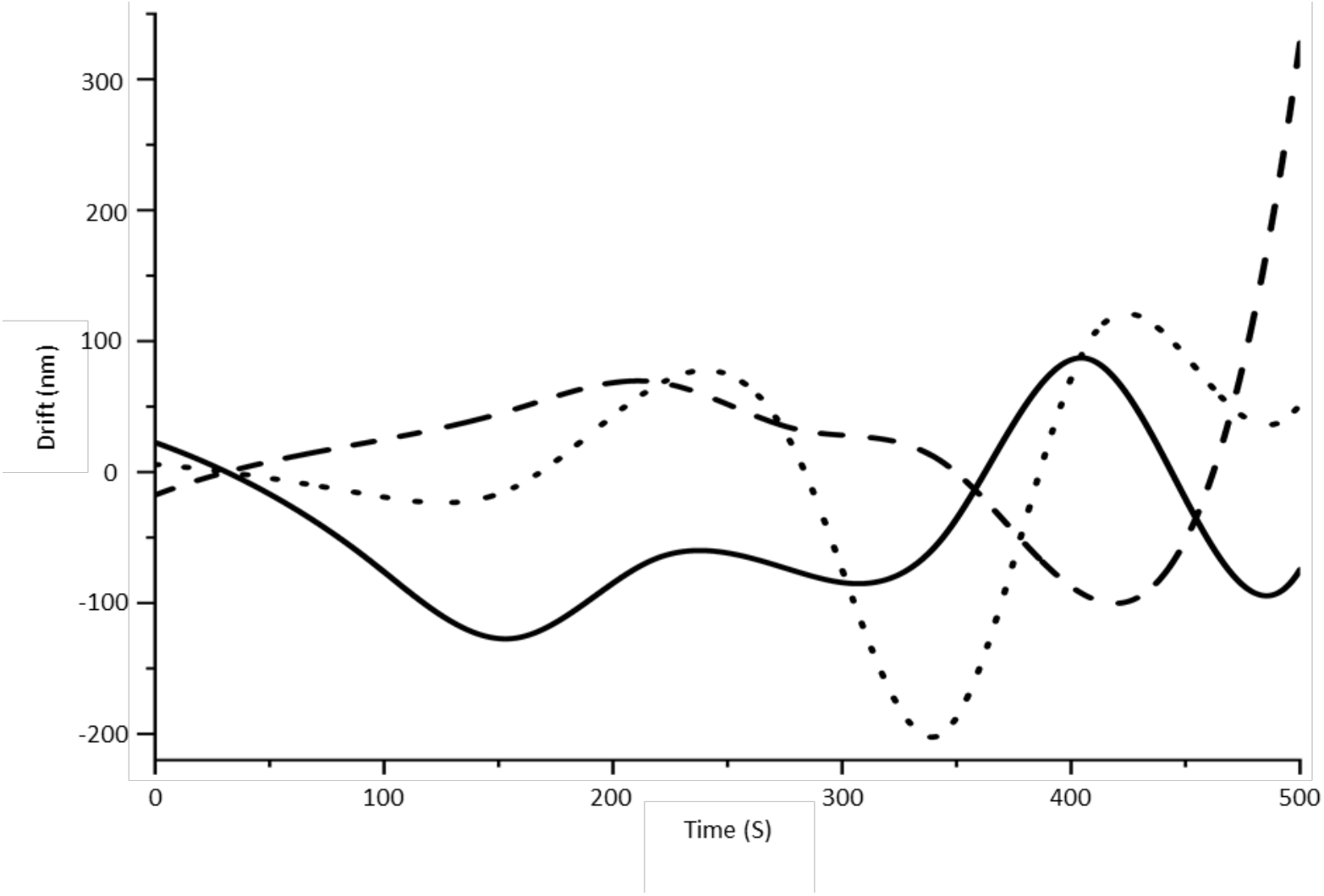
500 seconds movie recording using 50ms frame time and a 50nm diameter bead sample. To measure the drift in three dimensions, we used a saddle point PSF enabled by the deformable mirror. The figure shows the tracked position of the bead in X (straight line), Y (dotted line), and Z (dashed line) direction.

**Supplementary Figure 2.** We used phalloidin antibody conjugated with Alexa 647 for a control experiment to see whether these antibodies attach to phosvitins at droplets or not. Raw data were obtained by recording 10000 frames with 10 ms frame time. After analyzing the raw data and finding the localizations, we achieved a noisy super-resolved image (SFig2 A). A single frame from raw data is extracted to demonstrate, the blinking presents not only on the droplets but all over the field of view (SFig2 A).

**SFig 2.**
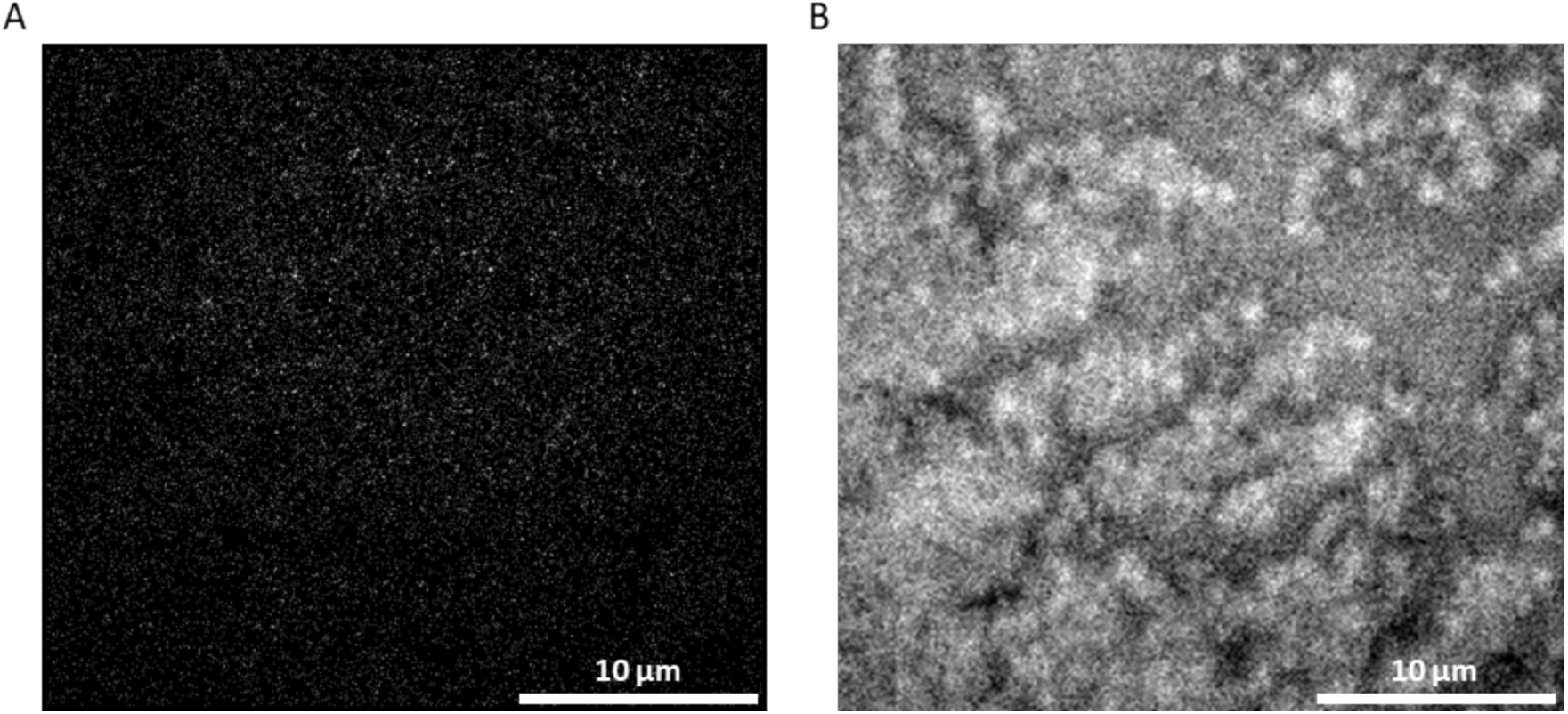
The control experiment using phalloidin antibody conjugated with Alexa 647. **A** *Super resolved image after processing the raw data from model emulsion with* phalloidin antibody. **B** A single frame from raw data to show the interaction of control antibody with droplets covered by phosvitins.

**Supplementary Figure 3.** To measure the apparent thickness of the oil droplets, a Gaussian was fitted to the line profiles of droplets in different imaging depths. For the imaging plane close to the surface, we measured a full width at half maximum of 71 nm with considering the average of both sides of the droplet (SFig3 A). With the same procedure we further fitted the apparent thickness of a droplet imaged in 4 μm depth (SFig3 B).

**SFig 3.**
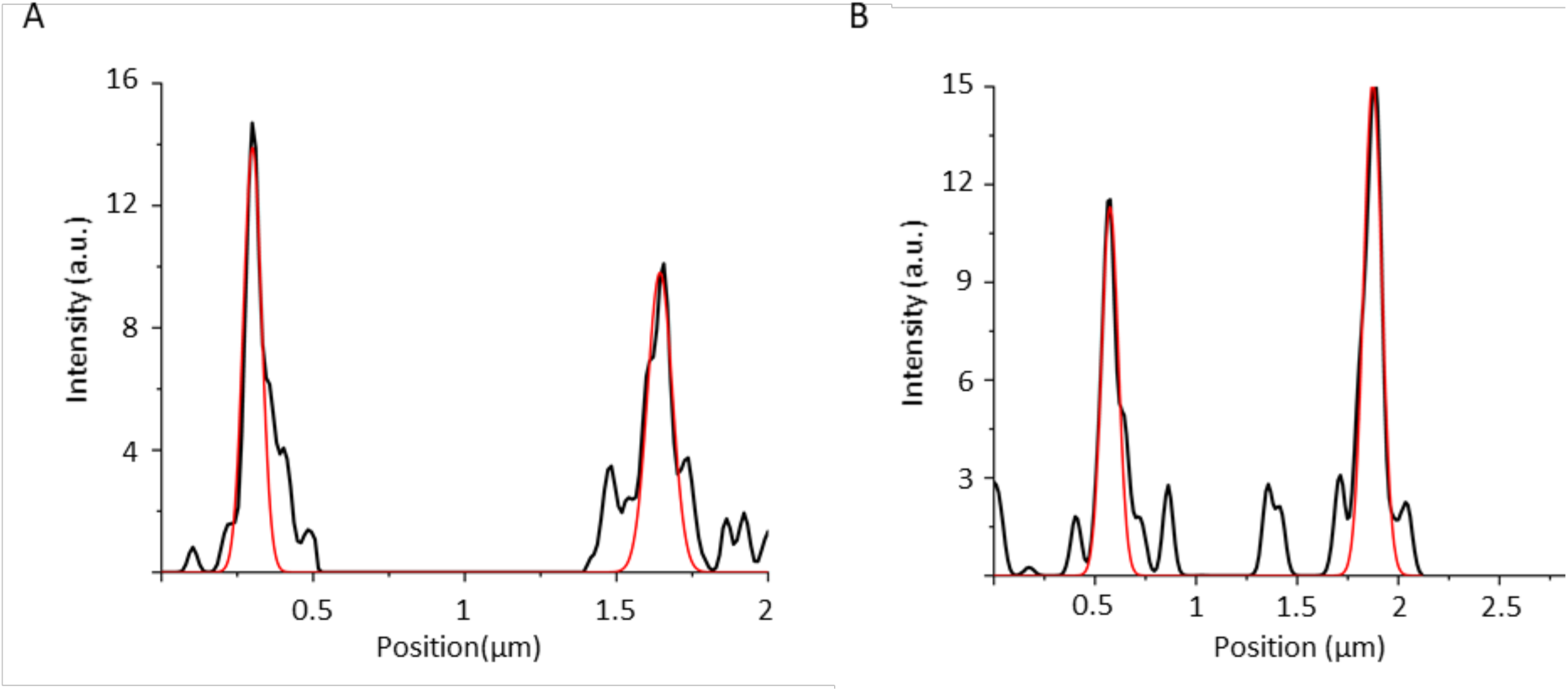
A Gaussian function was fitted to the intensity cross sections to calculate the apparent thickness of droplets in imaging different depths. **A)** Close to the surface. **B)** 4 μm in depth.

**Supplementary Figure 4.** To determine the resolution of super-resolved images from our model emulsion, we used Fourier ring correlation analysis. The localization list was transferred to the SMAP software to compute the resolution. The FRC curves show the decay of the correlation with spatial increasing frequency. After passing the threshold of 1/7, the resolution is calculated by taking the inverse of the spatial frequency at that point. For the droplets close to the surface, we determined a resolution of 71 nm (SFig4 A) and 124 nm for the imaging plane in 15 μm depth (SFig4 B).

**SFig 4.**
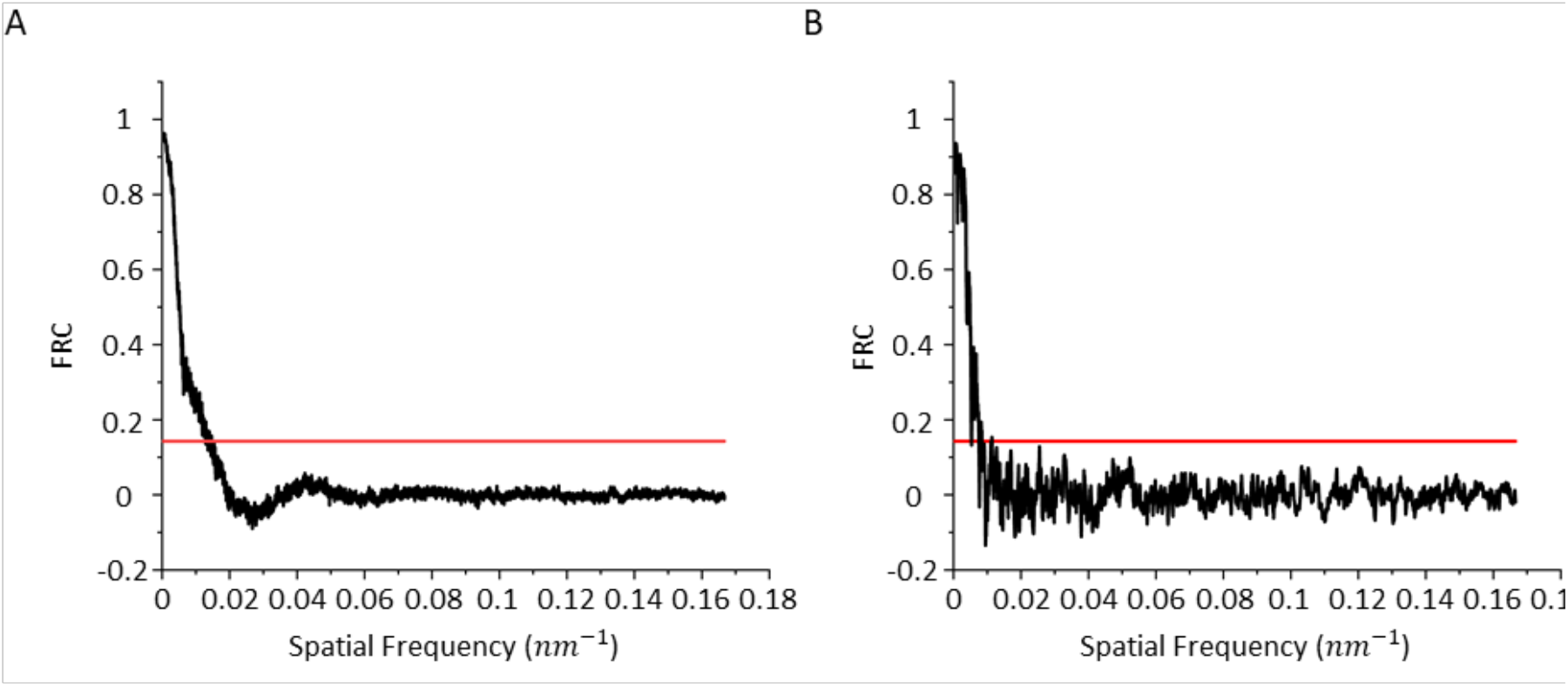
The data shows the FRC versus spatial frequency use to calculate the resolution of the images. **A)** close to the surface and, **B)** 4 μm in depth.

**Supplementary Figure 5.** Droplet distribution sizes were quantified using visual inspection and subsequent Hough circle transform. First, the droplets visually inspected based on the presence and absence of fluorescence. Consecutively, we used Hough circle transform for circles in the range between 0.2 μm to 2.0 μm to obtain the exact radius. Circles with radius lower than 0.2 μm and higher than 2.0 μm were removed from subsequent analysis. We used this approach for the image plane close to the surface (SFig5 A) and 4 μm in-depth (SFig5 B).

**SFig 5.**
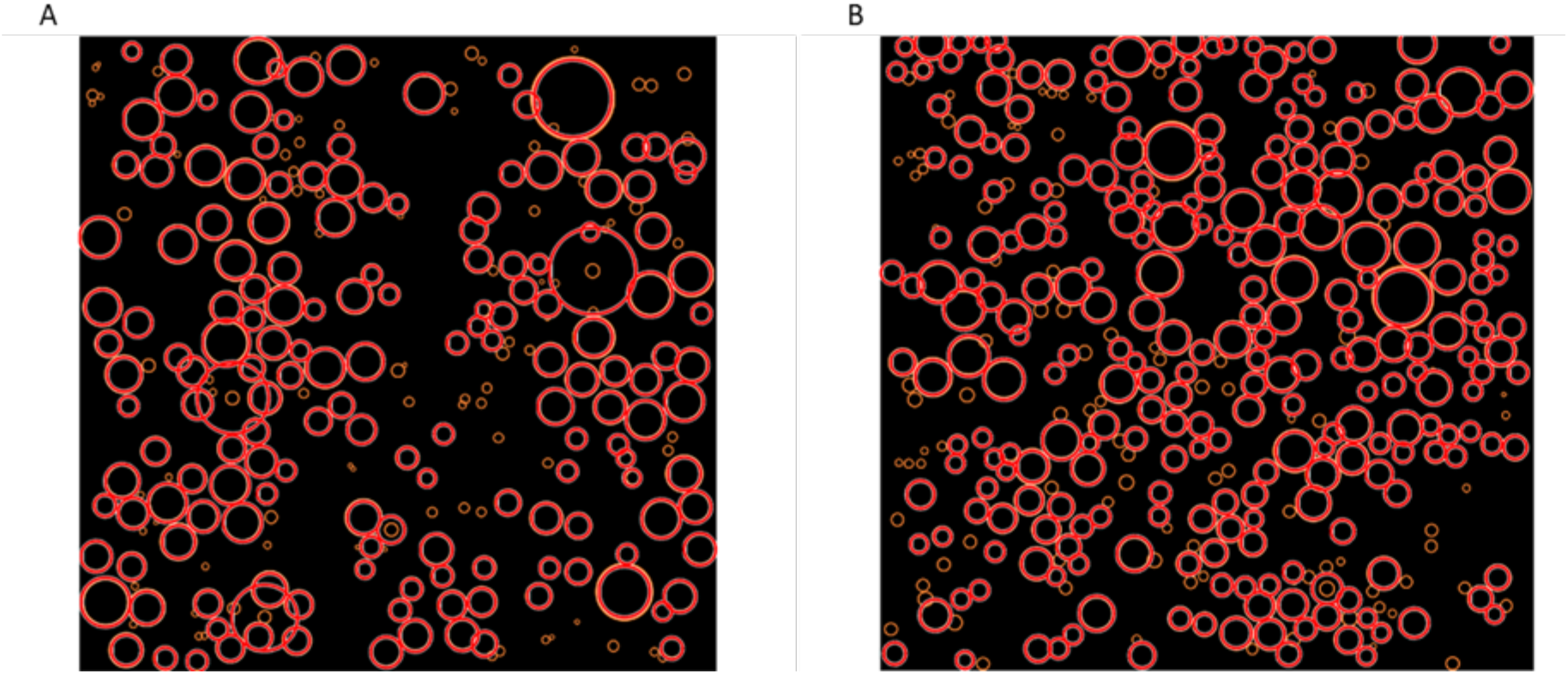
Analyzed field of view for droplet distribution. Orange circles show the visually inspected droplets and, red circles show detected droplets by Hough circle transformation for **A)** close to the surface and **B)** 4 μm in-depth.

